# Engineered human brown adipocyte microtissues improved glucose and insulin homeostasis in high fat diet-induced obese and diabetic mice

**DOI:** 10.1101/2021.10.11.463939

**Authors:** Ou Wang, Li Han, Haishuang Lin, Mingmei Tian, Shuyang Zhang, Bin Duan, Soonkyu Chung, Chi Zhang, Xiaojun Lian, Yong Wang, Yuguo Lei

**Author notes:** Corresponding Author Yuguo Lei, The Pennsylvania State University, PA, USA.

## Abstract

A large population of people is affected by obesity (OB) and its associated type 2 diabetes mellitus(T2DM). There are currently no safe and long-lasting anti-OB/T2DM therapies. Clinical data and preclinical transplantation studies show that transplanting metabolically active brown adipose tissue (BAT) is a promising approach to prevent and treat OB and its associated metabolic and cardiovascular diseases. However, most transplantation studies used mouse BAT, and it is uncertain whether the therapeutic effect would be applied to human BAT since human and mouse BATs have distinct differences. Here, we report the fabrication of three-dimensional (3D) human brown adipose microtissues, their survival and safety, and their capability to improve glucose and insulin homeostasis and manage body weight gain in high-fat diet (HFD)-induced OB and diabetic mice.

**Methods:** 3D BA microtissues were fabricated and transplanted into the kidney capsule of Rag1^-/-^ mice. HFD was initiated to induce OB 18 days after transplantation. A low dose of streptozotocin (STZ) was administrated after three month’s HFD to induce diabetes. The body weight, fat and lean mass, plasma glucose level, glucose tolerance and insulin sensitivity were recorded regularly. In addition, the levels of human and mouse adipokines in the serum were measured, and various tissues were harvested for histological and immunostaining analyses.

**Results:** We showed that 3D culture promoted BA differentiation and uncoupling protein-1 (UCP-1) protein expression, and the microtissue size significantly influenced the differentiation efficiency and UCP-1 protein level. The optimal microtissue diameter was about 100 µm. Engineered 3D BA microtissues survived for the long term with angiogenesis and innervation, alleviated body weight and fat gain, and significantly improved glucose tolerance and insulin sensitivity. They protected the endogenous BAT from whitening and reduced mouse white adipose tissue (WAT) hypertrophy and liver steatosis. In addition, the microtissues secreted soluble factors and modulated the expression of mouse adipokines. We also showed that scaling up the microtissue production could be achieved using the 3D suspension culture or a 3D thermoreversible hydrogel matrix. Further, these microtissues can be preserved at room temperature for 24 hours or be cryopreserved for the long term without significantly sacrificing cell viability.

**Conclusion:** Our study showed that 3D BA microtissues could be fabricated at large scales, cryopreserved for the long term, and delivered via injection. BAs in the microtissues had higher purity, and higher UCP-1 protein expression than BAs prepared via 2D culture. In addition, 3D BA microtissues had good in vivo survival and tissue integration, and had no uncontrolled tissue overgrowth. Furthermore, they showed good efficacy in preventing OB and T2DM with a very low dosage compared to literature studies. Thus, our results show engineered 3D BA microtissues are promising anti-OB/T2DM therapeutics. They have considerable advantages over dissociated BAs or BAPs for future clinical applications in terms of product scalability, storage, purity, quality, and in vivo safety, dosage, survival, integration, and efficacy.

## Introduction

According to World Health Organization data, about 10% of adults have obesity (OB), and the population with OB-associated T2DM will reach 300 million by 2025[1]. There are currently no safe and long-lasting approaches to prevent/treat OB and T2DM[2]. Healthy humans have a substantial amount of BAT, a tissue that augments the whole-body energy expenditure. Large clinical data shows BAT activity inversely correlates to body mass index, plasma glucose, and triglycerides levels, and insulin resistance, and BAT activity is a negative predictor of T2DM, dyslipidemia, coronary artery disease, cerebrovascular disease, congestive heart failure, and hypertension[3–5]. Furthermore, the beneficial effects of BAT are more pronounced in obese individuals, suggesting the importance of this tissue for the obese population[3–5].

Clinical studies have shown that augmenting BAT activities using pharmaceuticals (e.g., mirabegron[6], glucocorticoids[7], BIBO3304[8]), or cold stimulation[9–12] enhances the whole-body energy expenditure, glucose tolerance, and insulin sensitivity[13–15]. These findings suggest that BAT is a promising therapeutic target. However, these BAT-activating approaches require sustained treatments, have significant side effects[16], and may not work long-term. Additionally, they may work poorly on patients with a low abundance of BATs, such as obese and senior individuals (a population mostly affected by OB and its associated diseases).

An alternative approach to overcome these problems is to augment BAT mass and activity via tissue transplantation. Several groups transplanted BAT (0.1-0.2 g) from healthy mouse donors to diet-induced or genetically obese mice[17–20]. The transplantation significantly reduced plasma glucose, triglyceride, lipid levels, body weight gain, fat composition, and hepatic steatosis[19,20] while increasing the body energy expenditure, oxygen consumption rate (OCR), glucose homeostasis, and insulin sensitivity[21,22]. Transplanted BAT could directly burn fatty acids and glucose and dissipate the energy as heat via non-shivering thermogenesis (e.g., act as an energy sink)[15,16]. They also secreted soluble factors and exosomes that enhanced glucose uptake and energy expenditure in the heart, muscle, and WAT (e.g., act as an endocrine organ)[13,23–27]. Although few studies have used T2DM mice as recipients, research found that transplanting mouse BAT into STZ-induced diabetic mice prevented and reversed type 1 diabetes mellitus (T1DM)[21,28].

To date, most transplantation studies used mouse BATs, and it is uncertain whether these therapeutic effects, cellular and molecular mechanisms would be applied to human BAT since human and mouse BATs have distinct differences[29]. However, human BAT is located at deep organs, e.g., the supraclavicular, perirenal/adrenal, and paravertebral regions, and isolating sufficient human BAT for transplantation and research is challenging[30]. The Tseng group recently isolated and immortalized human brown adipocyte progenitors (BAPs) and found that these BAPs, when transplanted with Matrigel, could prevent/reverse diet-induced OB and metabolic disorders(Wang et al>, 2020). Interestingly, they also showed that human white preadipocytes activated to express endogenous UCP-1 had similar effects, indicating the importance of UCP-1 protein[31]. The study transplanted a large number (1.5-2×10^7^ cells per animal) of proliferating BAPs, which have potential tissue overgrowth risk. It also used Matrigel, an extracellular matrix (ECM) extracted from mouse tumors that contain hundreds of undefined components. Matrigel is not compatible with clinical applications.

To address these limitations, we explored if transplanting a low dose of mature BAs (differentiated from the above BAPs) in the absence of Matrigel could alleviate HFD-induced OB, metabolic symptoms, and T2DM in mice. When harvesting the fully differentiated BAs cultured in the conventional two-dimensional (2D) cell culture dishes, the trypsin-based dissociation killed a large percentage of cells. Additionally, the remaining BAs had a very low survival rate after transplantation. We thus proposed to prepare injectable 3D BA microtissues to overcome these problems. Here, we report the BA microtissue fabrication method, their survival, safety, and capability to improve glucose and insulin homeostasis and manage body weight gain in HFD-induced OB and diabetic mice.

## Methods

### 2D cell culture and differentiation

Immortalized BAPs are gifts from Dr. Tseng at Harvard University(Xue et al>, 2015). We followed published methods to culture BAPs[33,34]. Briefly, BAPs were cultured in Dulbecco’s Modified Eagle Medium (DMEM, HyClone, #SH30003.03) supplemented with 10% FBS (Atlanta biologicals, #S11150). When cells reached 80% confluence, they were passaged (1:3) with 0.25% trypsin-EDTA (Giboco, #25200056). To induce differentiation, BAPs were seeded at 0.5×10^4^ cells/cm^2^ and maintained in the growth medium to reach confluence. Then cells were cultured in differentiation medium I consisting EBM-2 (Lonza, #CC-3156), 0.1% FBS, 5 μM SB431542 (Selleckchem, #S1067), 25.5 μg/ml ascorbic acid (Sigma, #A89605G), 4 μg/ml hydrocortisone (Sigma, #H0396), 10 ng/ml Epidermal Growth Factor (Peprotech, # 100-15), 0.2 nM 3,3′,5-Triiodo-L-thyronine (T3, Sigma, #T2877), 170 nM insulin (Sigma, #I9287-5ML), 1 μM rosiglitazone (Sigma, #R2408), 0.5 mM 3-Isobutyl-1-methylxanthine (IBMX, Sigma, #I5879) and 0.25 μM dexamethasone (Sigma, #D4902) for 3 days, then in differentiation medium II (differentiation medium I without IBMX and dexamethasone) with medium change once a week. Mature BAs could be obtained 20–30 days after induction.

### Fabricating 3D BA microtissues in microwells

Aggrewells (Stemcell Technologies #34815, #34425) were pre-treated with anti-adherence rinsing solution (Stemcell Technologies, #07010) following the manufacturer’s instruction. Single BAPs were seeded into Aggrewells with differentiation medium I. Differentiation medium II was used after three days and refreshed once a week. Differentiated BA microtissues were collected by centrifugation at 100 g for 3 min.

### Fabricating 3D BA microtissues in shaking plates

Single BAPs were suspended in differentiation medium I in a low adhesion 6-well plate (Corning, #3471) shaking at 75 rpm. Detailed methods of culturing cells in shaking plates can be found in our previous publications[35–39]. Differentiation medium II was used after three days and refreshed once a week. The plate was tilted and placed in static for 5 min to settle down the microtissues to change medium. 90% of the exhausted medium was replaced with fresh medium. BA microtissues were collected by pipetting the medium up and down to suspend the microtissues and spinning at 100 g for 3 min.

### Fabricating 3D BA microtissues in thermoreversible hydrogels

Single BAPs were suspended in growth medium I in low adhesion 6-well plate shaking at 75 rpm overnight to form BAP spheres. The spheres were then mixed with 10% ice-cold PNIPAAm-PEG (Cosmo Bio, #MBG-PMW20-5005) solution dissolved in DMEM medium. The mixture was then cast on a tissue culture plate and incubated at 37 °C for 10 min to form a hydrogel before adding the pre-warmed differentiation medium I. Differentiation medium II was used after three days and refreshed once a week. To harvest BA microtissues, the medium was removed, and ice-cold DPBS (Life Technologies, #21600044) was added to dissolve the hydrogel for 5 min. Finally, BA microtissues were collected by spinning at 100 g for 3 min. Detailed methods of encapsulating cells in this thermoreversible hydrogel can be found in our previous publications[35,39–43].

### 3D microtissues transplantation

The animal experiments were conducted following the protocols approved by the University of Nebraska–Lincoln Animal Care and Use Committee. 6 mice were used for each study group. 6-week-old male B6.129S7-Rag1tm1Mom/J or Rag1 knock-out mice (Rag1^-/-^) were purchased from Jackson. Mice were transplanted with 1.25 million cells suspended in DPBS. DPBS was used as a sham control. Briefly, 3D microtissues were collected and transferred into a sample loading tip bent to have a U-shape. The tips were connected to a PE50 tube (BD Diagnostics system, #427516), and microtissues were slowly pushed into the center of the tube using a pipetman. The PE50 tube with microtissues was then bent into U-shape and placed into a microcentrifuge tube with both the PE50 tube ends kinked and facing up. The EP tube was centrifuged at 1000 rpm for 10 seconds to pack the microtissues tightly so that the medium was located to the two ends. The PE50 tube was connected to a loading tip again. Microtissues were slowly pushed to one end of the PE50 tube using a pipetman. This operation was to remove the medium so that no medium was injected into the kidney capsule. Otherwise, the medium would wash out the injected microtissues from the kidney capsule.

The right kidney of an anesthetized mouse was exposed. A small scratch on the flank of the kidney was made by a syringe 25-gauge needle, creating a nick in the kidney capsule. Saline was applied with a cotton swab to keep the kidney wet. The PE50 tube was inserted into the capsule to make a small pocket under the capsule. Microtissues were slowly pushed into the pocket. The PE50 tube was carefully retracted, and the nick was cauterized with low heat. After stopping bleeding, saline was applied, and the kidney was placed back. Details can be found in previous publications (Szot, Koudria and Bluestone, 2007; Jofra et al>, 2018). After 18 days of transplantation surgery, Rag1^-/-^ mice without BA transplantation were fed with NCD (Research diets, #D12450Ji) or HFD (Research diets, #D12492i) and labeled as Rag1^-/-^ NCD and Rag1^-/-^ HFD, respectively. Rag1^-/-^ mice transplanted with BA microtissues were fed with HFD and labeled Rag1^-/-^ HFD+BA. Wild-type (WT) mice without BA transplantation were fed with NCD or HFD and labeled as WT NCD and WT HFD.

### Immunocytochemistry

Cells cultured on 2D were fixed with 4% paraformaldehyde (PFA) at room temperature for 15 min, permeabilized with 0.25% Triton X-100 for 10 min and blocked with 5% donkey serum for 1 h before incubating with primary antibodies (Table S1) in DPBS + 0.25% Triton X-100 + 5% donkey serum at 4 °C overnight. After washing, secondary antibodies (Table S1) were added and incubated at room temperature for 2 h followed by incubating with 10 mM 4’,6-Diamidino-2-Phenylindole, Dihydrochloride (DAPI) for 10 min. Cells were washed with DPBS three times before imaging with a Fluorescent Microscopy (Zeiss, Germany). For 3D microtissues immunostaining, microtissues were fixed with 4% PFA at 4 °C overnight. 40 μm thick tissue sections were obtained via cryosection. The sections were washed with DPBS three times and stained as the 2D cell cultures.

For in vivo studies, mouse tissues were harvested and fixed with 4% PFA at 4 °C overnight. 5 μm thick sections were obtained via paraffin-embedding and section. The sections were deparaffined with xylene three times and rehydrated sequentially in 100%, 95%, 70%, 50% ethanol, and distilled water. For hematoxylin and eosin (H&E) staining, rehydrated sections were stained in Mayers Hematoxylin (Ricca Chemical Company, #3530-16) for 1 min, washed in distill water for 5 times, in DPBS once, and in distilled water 3 times with 1 min for each wash and stained in Eosin (Fisher, #SE23-500D) for 1 min. The sections were then dehydrated in 95% ethanol (3 times), 100% ethanol (2 times), and xylene (3 times) with 1 min for each wash before being mounted with coverslips. For immunostaining, heat-induced epitope retrieval was done on rehydrated tissue sections using the antigen retrieval buffer (Abcam, #ab93680) following the manufacturer’s instruction. The sections were stained as the 2D cell cultures.

### Flow Cytometry

Cell culture or microtissues were dissociated into single cells with 0.25% trypsin-EDTA. Single cells were fixed with 4% PFA and stained with primary antibodies at 4 °C overnight. After washing (three times) with 1% BSA in DPBS, secondary antibodies were added and incubated at room temperature for 2 h. Finally, cells were washed with 1% BSA in DPBS and analyzed using CytoFLEX LX (Beckman Coulter, USA). Isotype controls served for negative gating.

### Serum preparation

Blood samples were collected from the mouse tail. Cells were removed via centrifugation at 3000 g for 10 min at 4 °C. The supernatant was transferred into a new tube and centrifuged at 3000 g for 5 min at 4 °C to collect the serum.

### Mouse adipokine array and human obesity array

Mice were sacrificed. Blood was collected, and serum was isolated as described above. 100 µL serum was applied to the mouse adipokine array (R&D systems, #ARY-013) following the manufacturer’s instruction. 70 µL serum was applied to the human obesity array (RayBiotech, #QAH-ADI-3-2) following the manufacturer’s instructions.

### Glucose tolerance test (GTT)

Mice were fasted from the morning for 16 h. Glucose (0.75 g/kg body weight) was administrated via intraperitoneal injection. Blood samples were collected to measure the glucose level at 0 (baseline), 30, 60, 90, and 120 min after injection.

### Insulin tolerance test (ITT)

Mice were fasted overnight for 8 h. Insulin (Sigma, # I9278-5ML, 0.75 U/kg body weight) was administered via intraperitoneal injection. Blood samples were collected to measure the glucose level at 0 (baseline), 30, 60, and 90 min after injection.

### Body composition

Mouse fat and lean mass were analyzed by Minispec LF50 (Bruker, USA).

### Statistical Analysis

The data are presented as the mean ± SEM. Unpaired t-test and one-way analysis of variance (ANOVA) were used to compare two and more than two groups, respectively. Two-way ANOVA was used to compare mice metabolic curves and arrays. *P* < 0.05 was considered statistically significant.

## Results

### Fabricating 3D BA microtissues

The brown adipose progenitors (BAPs) used in this study were isolated from the superficial neck fat of a human subject and have been well characterized and documented[32]. To prepare 3D BA microtissues, single BAPs were placed in microwells (Aggrewells) in the growth medium. Cells interacted, gradually contracted, and formed compact spheroids after 24 h (**Figure S1A**). Like the 2D differentiation, we cultured the microtissues in differentiation medium I for 3 days and then in differentiation medium II with a medium refresh every 7 days (**Figure 1A**). Microtissues grew in size significantly during the differentiation (**Figure S1A**). For instance, microtissues with an initial diameter of 100 µm became 200 µm on day 17 (**Figure S1A**). The microtissue size growth can result from the cell number increase due to cell proliferation or the cell size increase, or both. Cell proliferation and size growth were observed during the differentiation of BAPs to BAs in 2D culture[33]. Immunostaining showed that most of the cells in the microtissues expressed the UCP-1 protein, a biomarker of mature and functional BAs (**Figure 1B**). UCP-1 is a mitochondria membrane protein that is critical for non-shivering thermogenesis. BAT dissipates energy in the form of heat via the UCP-1 activity[46,47]. To study if the microtissue size influences the differentiation efficiency, we prepared microtissues with a diameter of 100, 250, and 450 µm (initial diameter) and differentiated them. All microtissues grew in size significantly (**Figure S2**). While the 100 and 250 µm microtissues maintained spherical, the 450 µm microtissues gradually became non-spherical (**Figure S2D**). Cell death (**Figure S2D**, red arrows) and fusion between microtissues (**Figure S2D**, blue arrows) were observed only in the 450 µm microtissues. On day 17, flow cytometry analysis showed 92.6%, 72.6%, and 80.4% of the cells in the 100, 250, and 450 µm microtissues were UCP-1 positive. For comparison, 2D culture resulted in 78.8% UCP-1 positive cells (**Figure 1C**). The mean fluorescent intensity (MFI) of UCP-1 intensity in 100 µm microtissues is significantly higher than in other groups (**Figure 1D**). The results show that a 3D microenvironment promoted brown adipogenesis and UCP-1 expression. However, a small microtissue size should be used to maximize the benefits. We used 100 µm microtissues for the rest of the study.

**Figure 1.**
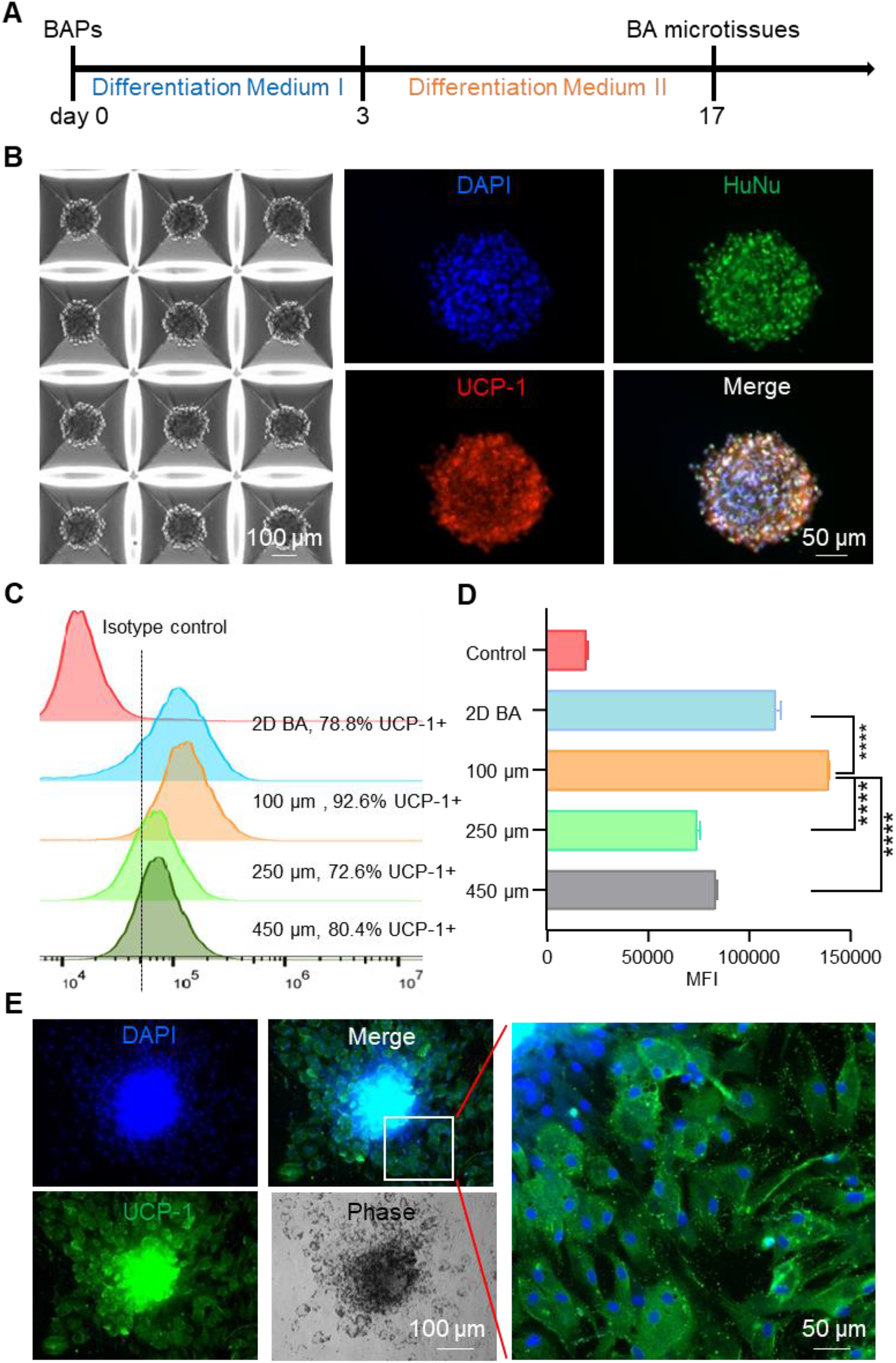
3D culture enhanced BA differentiation. (**A**) BA differentiation protocol. (**B**) 3D BA microtissues in microwells on day 17 and their immunostaining. HuNu: human nuclear antigen. (**C**) Flow cytometry analysis of UCP-1 expression on day 17 for BAs prepared in 2D culture and 3D culture with varied aggregate sizes. (**D**) The mean fluorescent intensity (MFI) of UCP-1 as measured with flow cytometry in (**C**). (**E**) The day 17 BA microtissues were plated on 2D surface for 6 days and stained for UCP-1 expression. Data are represented as mean ± SEM (n=3). ****p < 0.0001.

When the day 17 microtissues were placed in a conventional tissue culture plate, they adhered to the surface, and individual cells migrated out (**Figure 1E, S1B, S1C**). These cells had the classical BA characteristics. They expressed a high level of UCP-1 protein (**Figure 1C, 1D, 1E, S3D**) and had multiple small oil droplets (**Figure 1E, S3A, S3B, S3C**) and a high mitochondrial content (**Figure S3C**). In addition, they expressed the typical BA marker genes at high levels (**Figure S3E**).

### BA microtissues survived in vivo with angiogenesis and innervation

As said, very few BAT transplantation studies used T2DM mice. To fill the gap, we used high fat diet (HFD)-induced OB and T2DM mouse model to test the engineered BA microtissues. We selected the immune-deficient Rag1^-/-^ mice as recipients since they tolerate xenogeneic and allogeneic tissues and have HFD-induced OB and metabolic disorders[48,49]. In addition, they developed diabetes when given STZ[50]. Rag1^-/-^ mice fed with normal chow diet (NCD) and HFD with no microtissue transplantation were used as positive and negative controls to evaluate the microtissue efficacy. Wild type (WT) mice fed with NCD and HFD were also included to assess the difference of response to HFD between WT and Rag1^-/-^ mice. HFD was initiated 18 days post-transplantation (**Figure 2A**). A significant number of adipose tissue was found in the kidney capsule 1 month and 5 months after transplantation (**Figure 2B**). H&E staining showed the dense kidney tissue and the adjacent adipose tissue with large amounts of oil droplets (**Figure 2B**). The oil droplets were bigger in the 5 month sample. No tumor or non-adipose tissues were found in the transplants, indicating safety of fully differentiated BAs in vivo.

**Figure 2.**
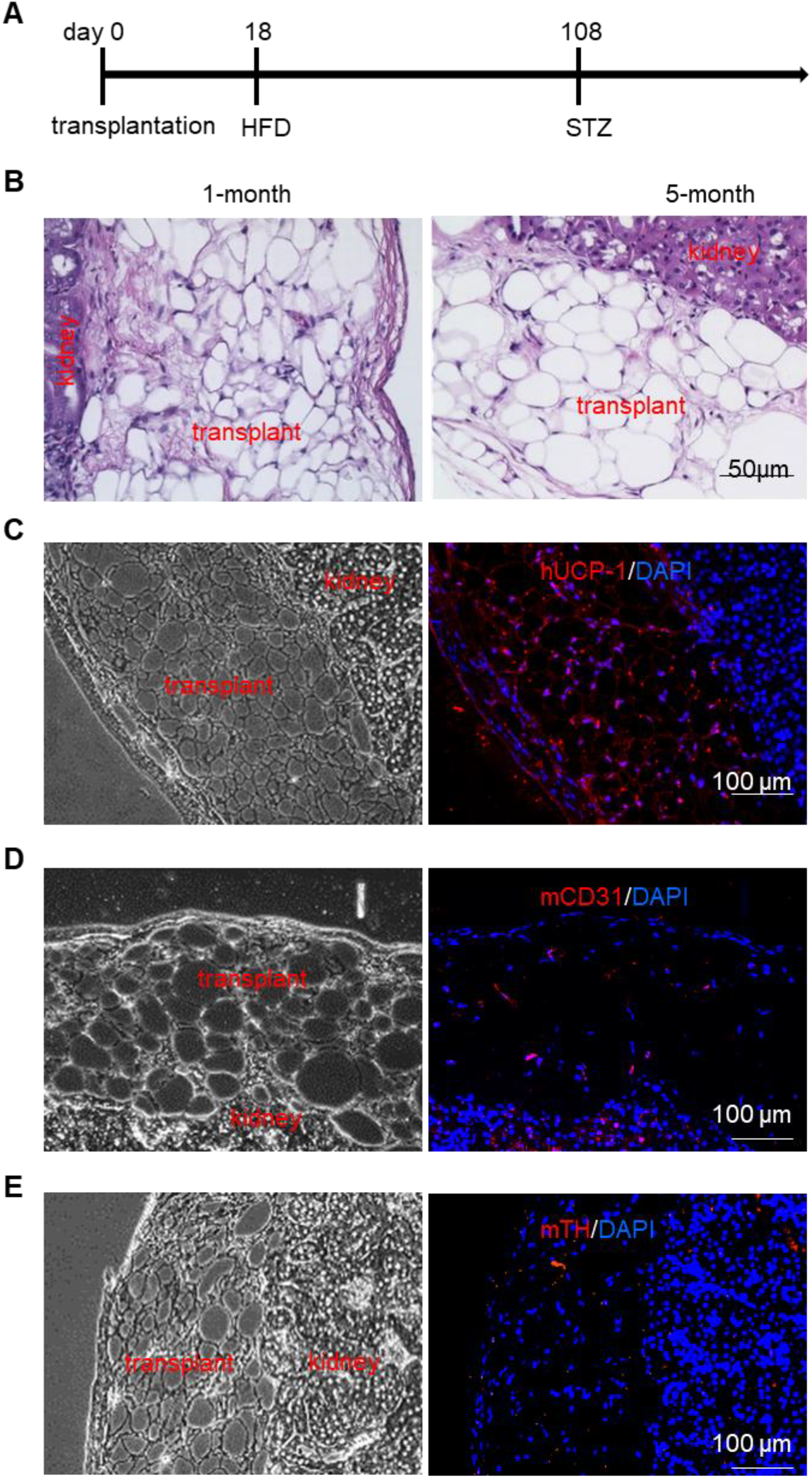
BA microtissues survived in vivo with angiogenesis and innervation. (**A**) The transplantation protocol. (**B**) H&E staining of 1-month and 5-month transplant. The transplant and adjacent kidney tissue are labelled. Immunostaining of the 5-month transplant and adjacent kidney tissue with human UCP1 (hUCP1) (**C**), mouse CD31 (mCD31) (**D**), and mouse tyrosine hydroxylase (mTH) (**E**).

The oil droplets were much bigger than those of the endogenous BAT (endoBAT) of NCD fed mice, but comparable to the endoBAT of HFD fed mice (**Figure 4A**), which indicates whitening of the transplanted BAs. However, a high level of UCP-1 protein was observed (**Figure 2C**), showing the BA function was maintained for at least 5 months. Host blood vessels grew into the transplants (**Figure 2D**). The vessel density was lower than the endoBAT of NCD mice but comparable to the endoBAT of HFD mice (**Figure 4D**). Innervation is critical for BA function. Tyrosine hydroxylase (TH+) nerves were found in the transplants (**Figure 2E**), and their density was lower than this of the NCD mouse endoBAT but comparable to the density in the HFD mouse endoBAT **(Figure 4E**).

**Figure 3.**
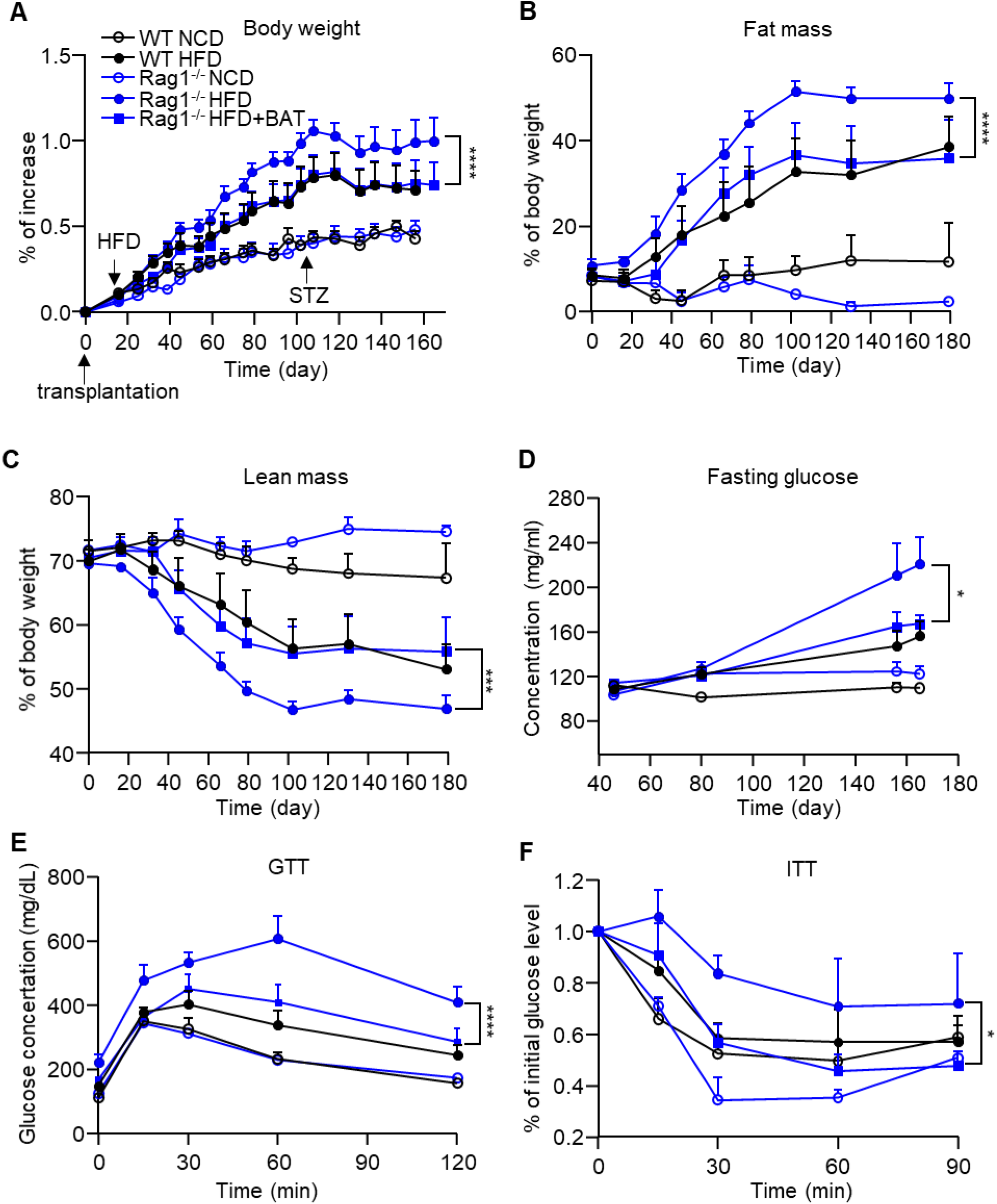
BA microtissues alleviated obesity and diabetes. (**A**) Body weight gain, (**B**) % fat mass, (**C**) % lean mass, (**D**) fasting glucose level, (**E**) GTT (day 150), and (**F**) ITT (day 170). WT: wild type mouse; Rag1^-/-^: Rag1 knock-out mice; NCD: normal chow diet; HFD: high fat diet. Data are represented as mean ± SEM (n=6). *p < 0.05, ***p < 0.001, ****p < 0.0001.

**Figure 4.**
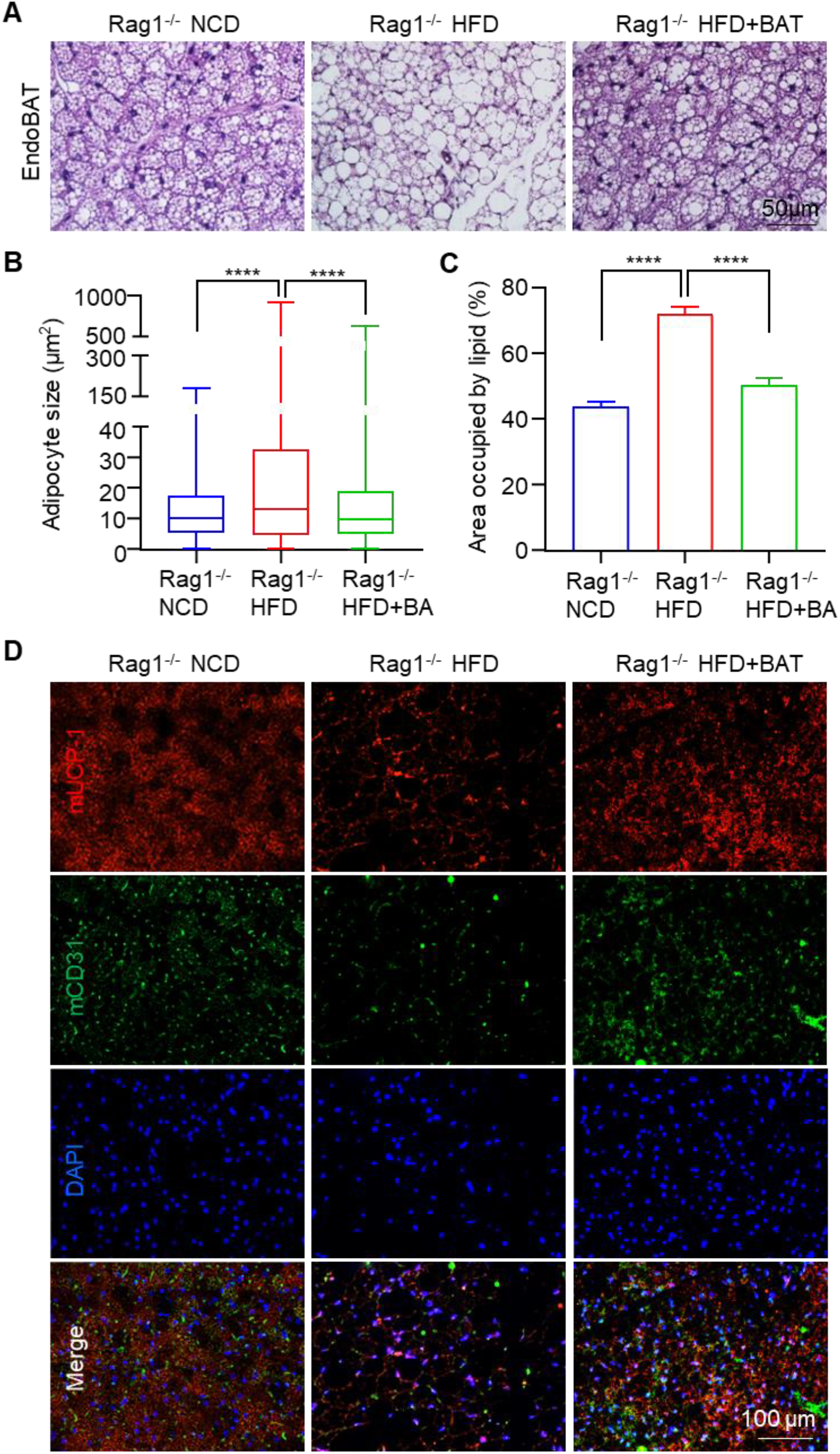
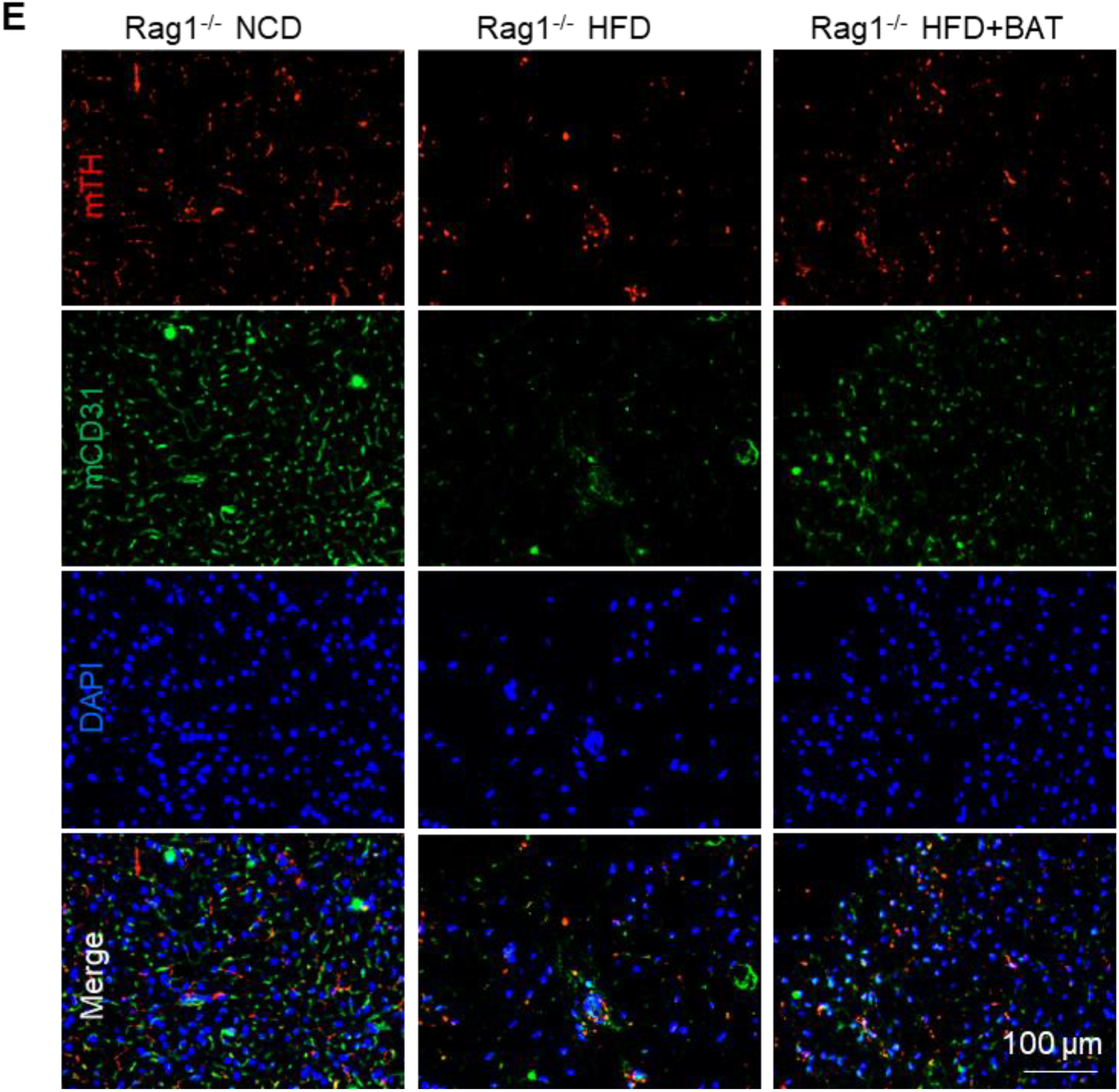
BA microtissues protected endogenous BAT (endoBAT). (**A**) H&E staining of endoBAT. BA size (**B**) and area (**C**) in endoBAT. (**D**) Mouse UCP-1 and CD31 expression in endoBAT. (**E**) Mouse TH and CD31 expression in endoBAT. Data are represented as mean ± SEM (n=6). ****p < 0.0001.

### BA microtissues alleviated obesity and diabetes

We regularly measured the body weight, fat and lean mass, plasma glucose level, glucose tolerance, and insulin sensitivity (**Figure 3**). To mimic the β cell dysfunction of T2DM, we injected low-dose STZ (90 mg/kg) after 3-month HFD. The bodyweight drops and the fasting glucose level rise after the STZ injection indicated the progression of T2DM (**Figure 3A, 3D**). There was no significant difference in total diet intake between the HFD groups. Under HFD, both WT and Rag1^-/-^ mice developed OB (**Figure 3A**) and grew large fat masses (**Figure 3B, 3C**). We found Rag1^-/-^ mice were slightly more prone to HFD-induced OB and metabolic disorders but were suitable for our studies. The transplant significantly reduced the body weight gain, fat content, and fasting glucose level while increasing insulin sensitivity and glucose tolerance (**Figure 3**).

### BA microtissues prevented the whitening of endogenous BATs

The endoBAT was whitened in Rag1^-/-^ mice fed with HFD, as shown by an increase of adipocyte size and oil droplets (**Figure 4A-C**), and a reduction of the mUCP-1 protein expression (**Figure 4D**), blood vessel density (**Figure 4D, 4E**) and TH+ nerve density (**Figure 4E**). The transplanted BA microtissues almost wholly prevented the whitening of endoBAT (**Figure 4A-E**). HFD mice with BA microtissues had similar BA morphology, blood vessel, and nerve densities to the NCD mice.

### BA microtissues alleviated endogenous WAT hypertrophy and liver steatosis

The adipocyte and oil droplet size of subcutaneous WAT (scWAT) was increased by the HFD. BA microtissues significantly reduced the adipocyte and oil droplet size (**Figure 5A-C**). CD31 staining showed fewer blood vessels in HFD fed mice. BA microtissues increased the blood vessel density in HFD fed mice (**Figure 5D**). The liver fat content was significantly increased in HFD mice but was almost normalized by BA microtissues (**Figure 5E**). These results showed that the transplanted microtissues had a profound effect on multiple organs.

**Figure 5.**
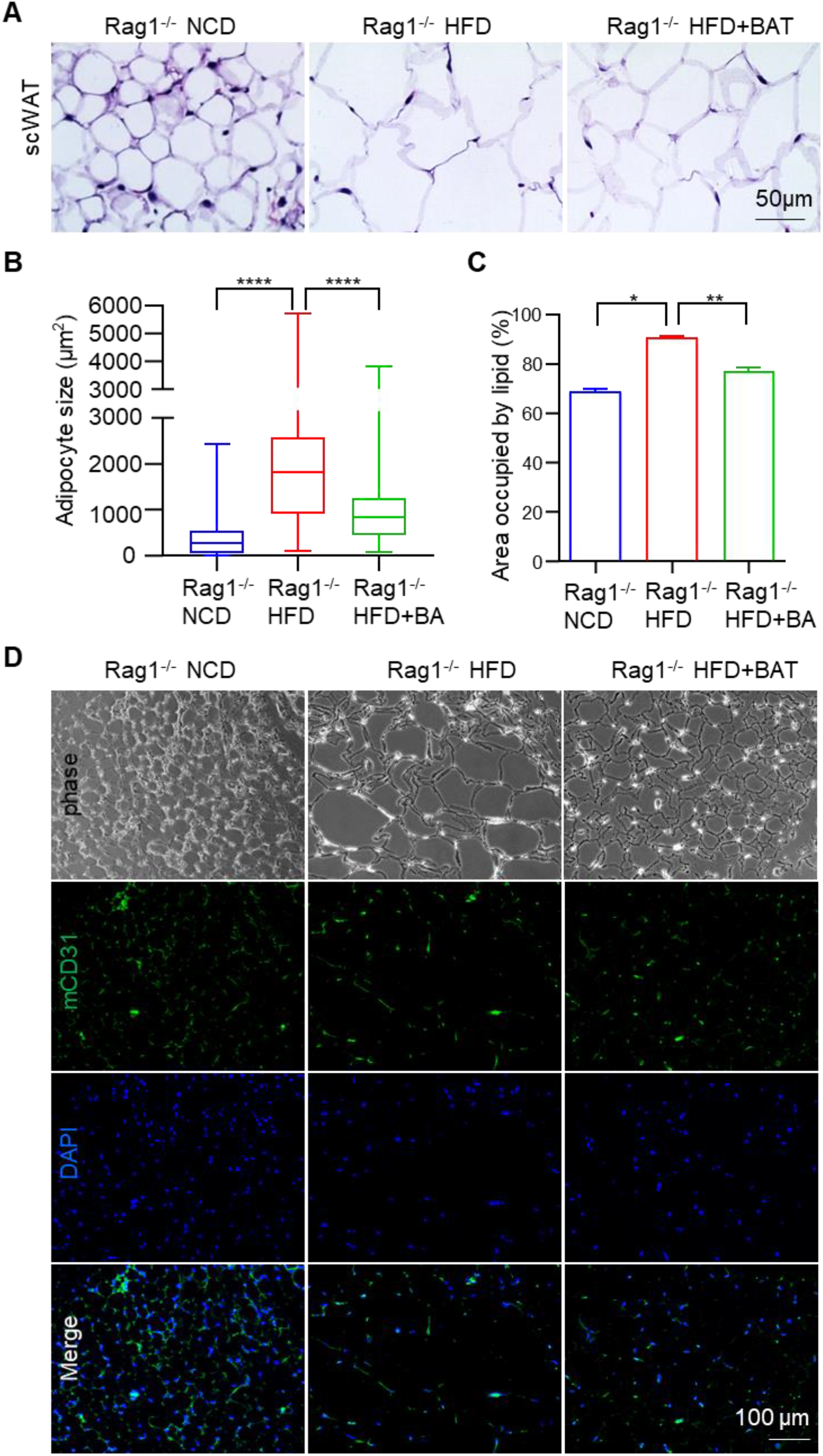
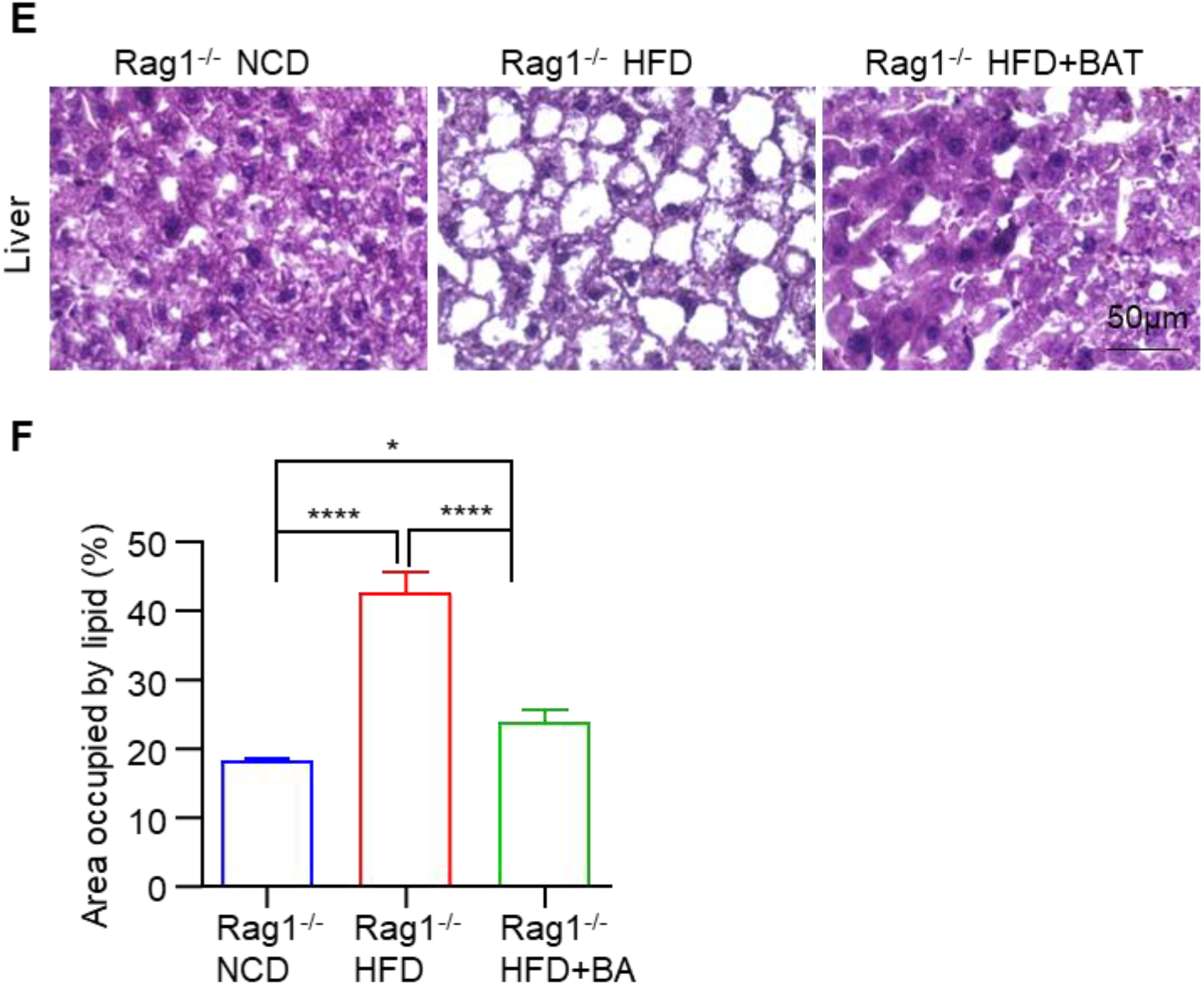
BA microtissues reduced endogenous WAT hypertrophy and liver steatosis. (**A**) H&E staining of mouse subcutaneous white adipose tissue (scWAT). (**B, C**) Adipose size and area in scWAT. (**D**) Mouse CD31 expression in scWAT. (**E**) H&E staining of mouse liver. (**F**) Adipose area in liver. Data are represented as mean ± SEM (n=6). *p < 0.05, **p < 0.01, ****p < 0.0001.

### BA microtissues secreted soluble factors and modulated endogenous adipokines

We used a Human Obesity Antibody Array to measure human protein factors in the blood. We found that the concentrations of human adiponectin, chemerin, and TGF-β1 in mice with transplants were significantly higher than background (**Figure 6A**), indicating the transplanted BAs were secreting soluble factors. Research has shown that adiponectin secreted by white and brown adipocytes has a protective role in obesity-associated metabolic and cardiovascular diseases. Adiponectin influences multiple tissues such as the liver, skeletal muscle, and vascular system. Adiponectin can increase insulin sensitivity[51,52], suppress inflammation, and reduce atherosclerosis[53–55]. Plasma adiponectin level is negatively correlated with obesity, and adiponectin deficiency enhances HFD induced insulin resistance[56]. Chemerin plays a positive role in the metabolic health[57,58]. TGF-β1 is a mediator of insulin resistance in metabolic disease[59,60].

**Figure 6.**
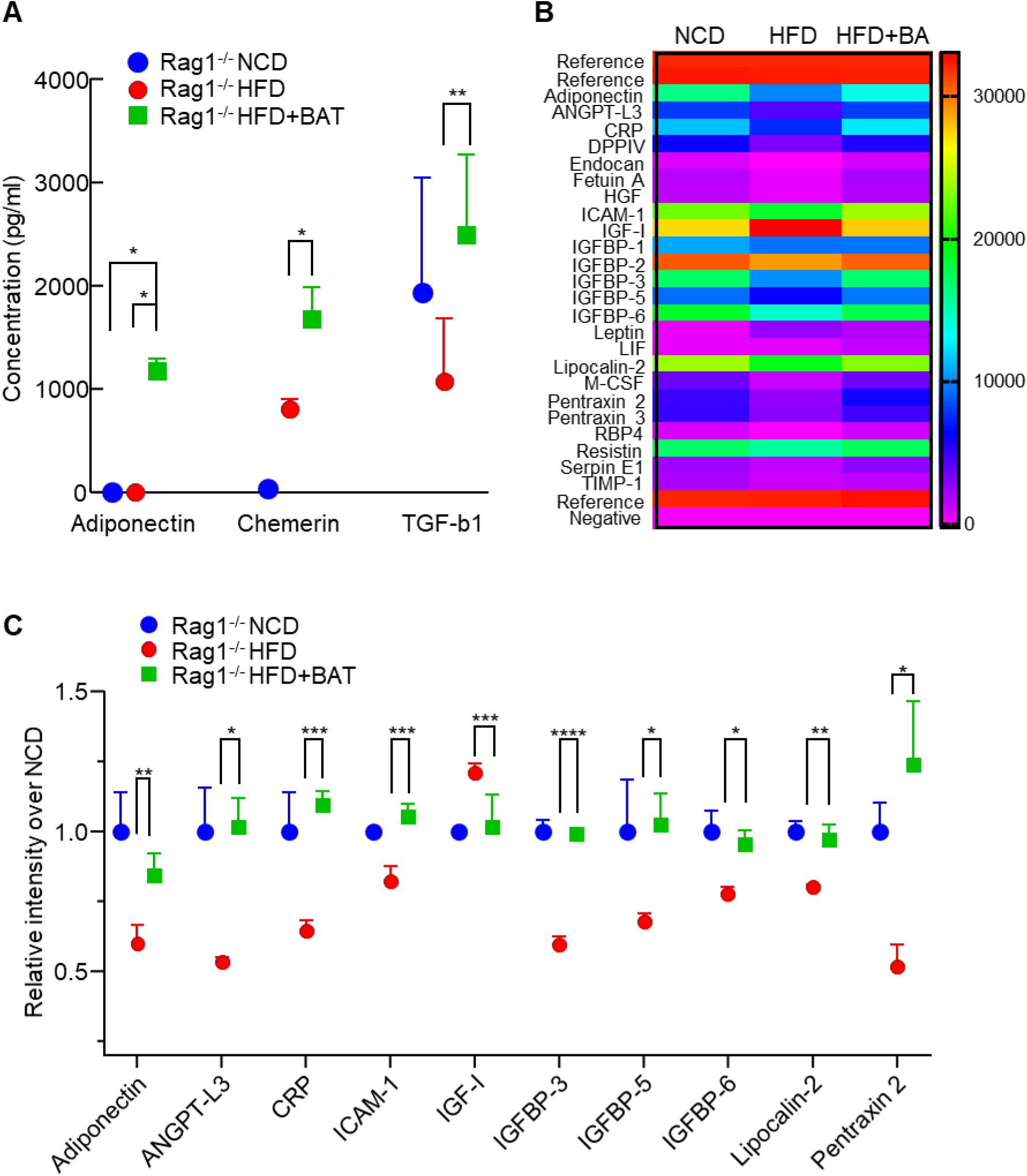
BA microtissues secreted human protein factors and modulated endogenous adipokines. (**A**) Protein factors in mouse plasma detected using human obesity antibody array. (**B, C**) Heatmap and quantification of adipokines in mouse plasma detected using mouse adipokine antibody array. Data are represented as mean ± SEM (n=3). *p < 0.05, **p < 0.01, ***p < 0.001, ****p < 0.0001.

We also used a Mouse Adipokine Antibody Array to measure 38 obesity-related mouse molecules. The levels of Adiponectin, ANGPT-L3, C-reactive protein, ICAM-1, IGFBP-3, IGFBP-5, IGFBP-6, Lipocalin-2, and Pentraxion2 were significantly reduced, and the IGF-1 concentration was increased by the HFD (**Figure 6B, 6C**). All of these molecules were normalized by the transplanted BA microtissues (**Figure 6C**). Thus, the transplanted microtissues had a significant effect on the endogenous adipokines.

### Scale up BA microtissues production

All the above experiments used microwells to prepare microtissues. However, microwells are unsuitable for producing microtissues at large scales, which are needed for future extensive animal studies, drug screens, and clinical applications. Previously, our lab used 3D suspension culture (e.g., shaking plates, spinning flasks, stirred tank bioreactors) to prepare human pluripotent stem cells at large scales[35–39]. We also pioneered in culturing stem cells at high density in thermoreversible PNIPAAm-PEG hydrogels[35,39– 43,61]. We found that BA microtissues could be prepared in both systems (**Figure 7**). Large tissue aggregates formed via fusion of multiple microtissues were found in shaking flasks (**Figure 7A**), while microtissues in PNIPAAm-PEG hydrogels were uniform (**Figure 7B**). BAs from both systems expressed a high level of UCP-1 protein (**Figure 7C)**. Thus, both systems can be used to scale up the microtissues production.

**Figure 7.**
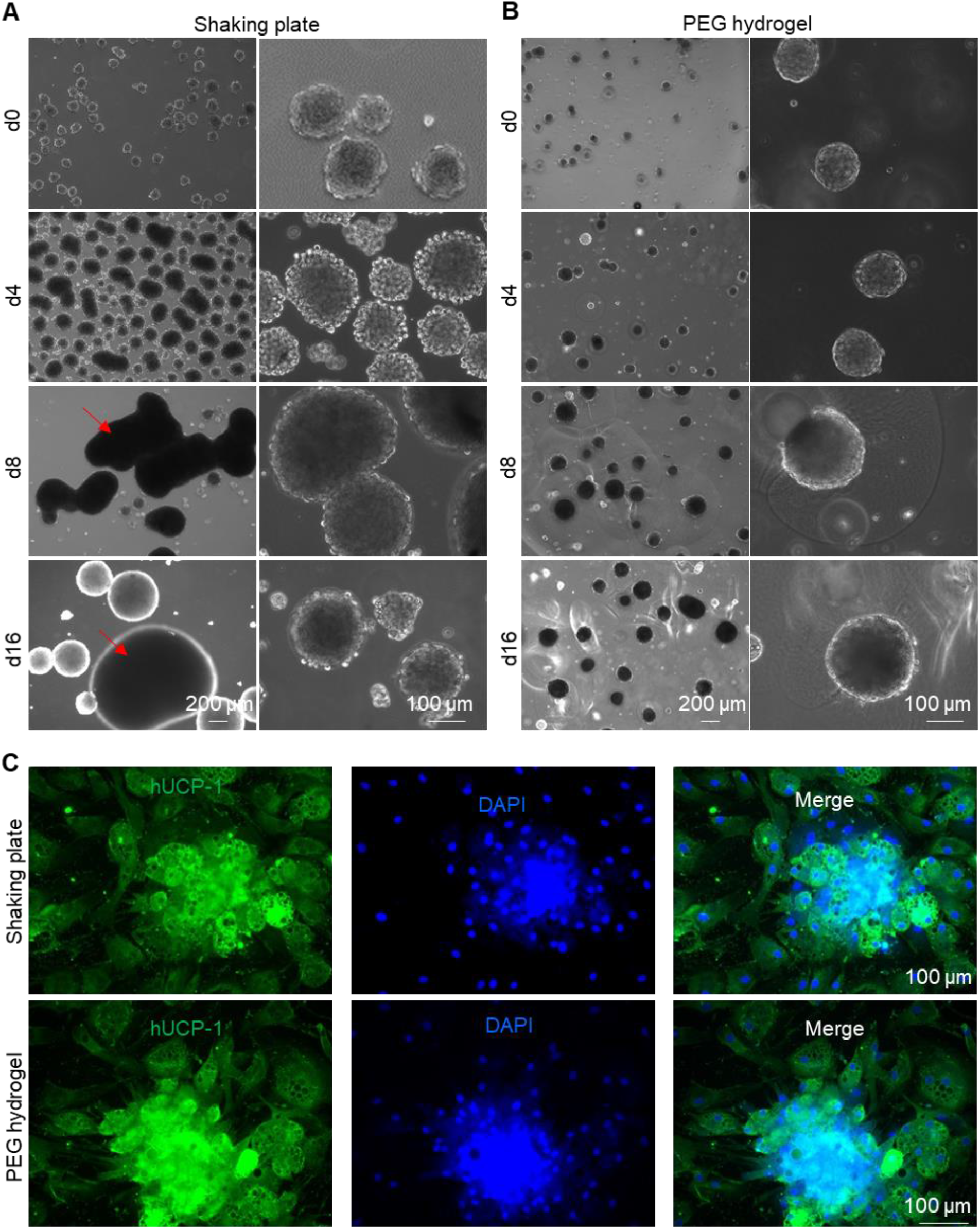
Scaling up BA microtissue production. Preparing BA microtissues in shaking plates (**A**) and PEG hydrogel (**B**). (**C**)Immunostaining of day 17 BA microtissues prepared in shaking plates and PEG hydrogel. Microtissues were plated on 2D surface for 6 days before staining.

### Preserving BA microtissues

Lastly, we evaluated if BA microtissues could be preserved. Microtissues preserved in cell culture medium at room temperature for 24 h (**Figure 8B**) had similar viability to the fresh sample (**Figure 8A**). Microtissues could also be cryopreserved in liquid nitrogen for the long term without significantly sacrificing cell viability (**Figure 8C**).

**Figure 8.**
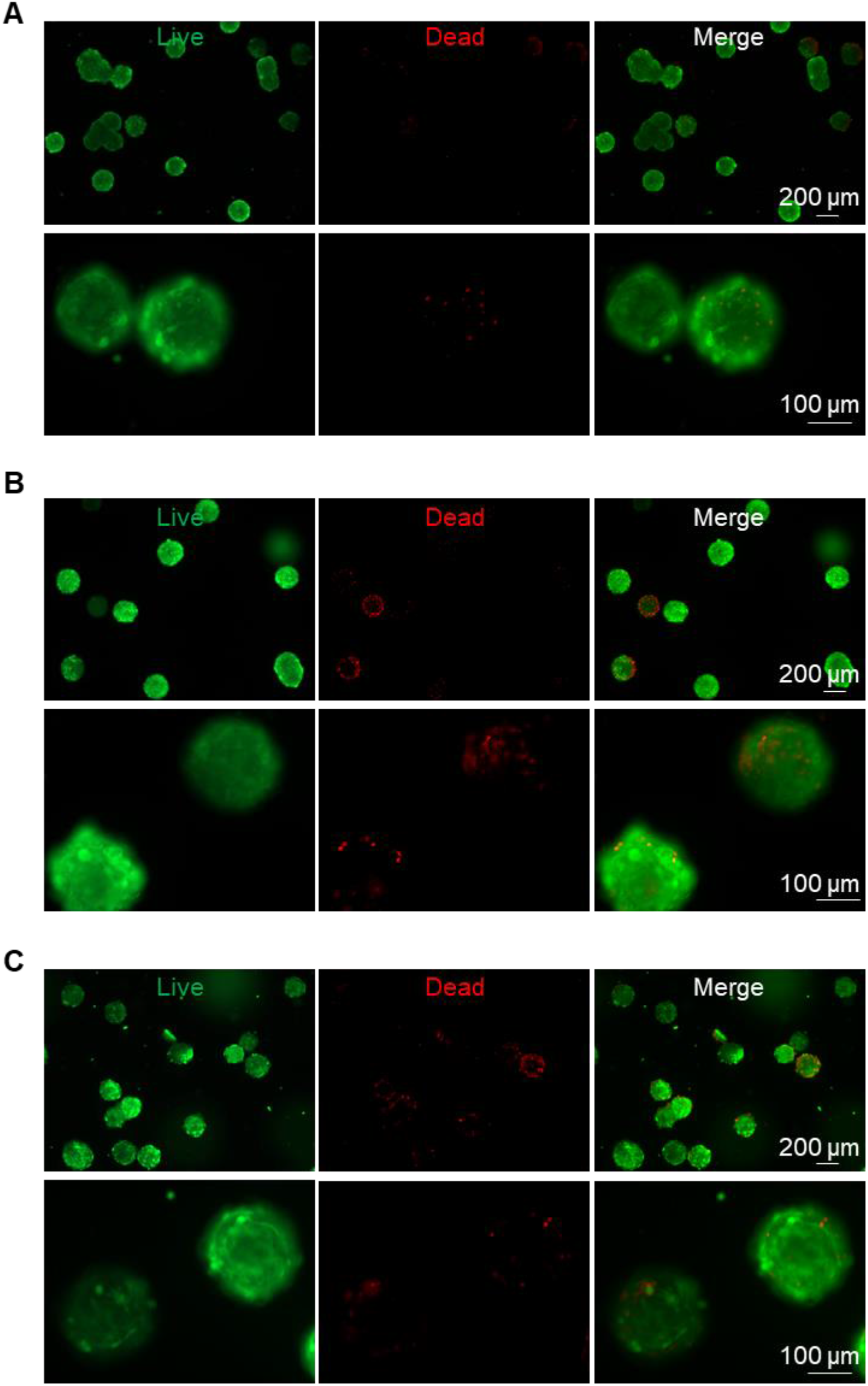
Preserving BA microtissues. Live/dead cell staining of day 27 fresh BA microtissues (**A**), or stored at room temperature for 24 h (**B**), or stored in liquid nitrogen and recovered (**C**).

## Discussion

Our study showed that human BAPs could be differentiated into mature BAs in 3D to prepare injectable BA microtissues (**Figure 1, S1, S2, S3**). The 3D culture promoted BA differentiation and UCP-1 protein expression. BA microtissues could survive in vivo for the long term with angiogenesis and innervation (**Figure 2**). They alleviated the body weight and fat gain and improved glucose tolerance and insulin sensitivity significantly in HFD-induced OB and diabetic mice (**Figure 3**). The transplanted BA microtissues impacted multiple tissues such as endogenous BAT, WAT, and liver (**Figure 4, 5**). In addition, they secreted protein factors and influenced the secretion of endogenous adipokines (**Figure 6**). These microtissues could be produced using the scalable 3D suspension culture or in a PEG hydrogel (**Figure 7**) and could be cryopreserved and shipped at room temperature (**Figure 8**). To our best knowledge, this is the first report on engineering human BA microtissues and showing their safety and efficacy in OB and T2DM mice. The findings that 3D culture promotes BA differentiation and the microtissue size affects the differentiation are all new.

To date, most BAT transplantation studies used mouse BATs. There is a need to study if human BAT can manage OB and its associated metabolic disorders. The Tseng group isolated and immortalized human BAPs and demonstrated that these cells could differentiate into mature BAs in vivo and reduce the HFD-induced OB and metabolic symptoms[31,32]. They transplanted proliferating BAPs instead of mature BAs that have exited the cell cycle. The possible reason is that BAPs have a better survival rate in vivo. A recent study showed that about 12.1% of immortalized mouse BAPs survived in SCID mice 7 days after transplantation with Matrigel, while only 2.7% of mature BA (differentiated from BAPs in vitro) were live after 7 days using the same transplantation procedure[18]. Consequently, only BAPs showed an efficacy in vivo[18]. However, there are potential problems with using BAPs. First, the differentiation efficiency in vivo is typically low[18]. Second, proliferating cells have a higher tumorigenic risk, especially for immortalized cells. Transplanting fully differentiated BAs has advantages in that they can be prepared in vitro at high purity (e.g., ∼93% in this study) (**Figure 1**). They are less likely to have uncontrolled growth in vivo. However, approaches must be developed to improve their survival rate in vivo and avoid cell death during the harvest in vitro, as found in our preliminary studies.

Prior studies showed transplanting mouse BATs prevented/reversed OB and its associated metabolic disorders. It should be noted that only transplanting intact 3D BAT, not dissociated single BAs, achieved long-term survival and function[17–20], indicating that 3D BAT, not single BAs, should be used as therapeutics in the future. Researchers typically inject cultured BAPs with Matrigel to mimic a 3D tissue. Matrigel can both restrict the cells at the transplantation size and enhance their survival[18]. However, Matrigel is extracted from mouse tumor tissue and is not chemically defined, and not compatible with clinical applications. Our preliminary study showed that a large percentage of mature BAs died during the harvest process, suggesting that single BAs are unsuitable for transplantation. Our engineered BA microtissues are injectable, do not need extra matrix and dissociation, thus addressing all these problems.

Interestingly, we found that the microtissue size influenced the BA differentiation efficiency significantly. The optimal diameter is about 100 µm (initial diameter). BAs have a high demand for glucose, oxygen, and nutrients to meet their high metabolic activities[18,62–66]. Nutrients are transported in these microtissues mainly through diffusion. Therefore, cells at the center of large microtissues may not have a sufficient supply of nutrients, negatively affecting the differentiation process. A second possible cause is that cells secrete autocrine or/and paracrine factors are important to BA differentiation. The microtissue diameter influences the concentrations and gradients of these factors. The exact reason should be made clear in future studies.

We demonstrated that the production of BA microtissues could be scaled up using the 3D suspension culture or a 3D thermoreversible hydrogel matrix. A limitation with the 3D suspension culturing is that the microtissues are heterogeneous in size (**Figure 7A**). Using the thermoreversible hydrogel matrix can produce homogenous microtissues in size (**Figure 7B**). However, the matrix is expensive. Our group recently developed a novel microbioreactor termed AlgTubes, in which cells are cultured in alginate hydrogel microtubes[36,37,67–72]. AlgTubes are scalable and have low cost. The cell aggregate size can be precisely controlled by the hydrogel tubes. AlgTubes can produce up to 5×10^8^ cells per mL of culture volume, about 200 times more than the 3D suspension culture. Future studies can apply AlgTubes to produce fibrous BA microtissues with uniform and precise size at high density.

BAT is among the most vascularized tissues in the body, averaging ∼1.2 capillaries per BA (versus only ∼0.4 capillaries per white adipocyte) [62–66]. A substantial blood supply is required to provide glucose, fatty acids, nutrients, and oxygen to fuel thermogenesis and rapidly distribute heat throughout the body [73]. Vascular cells also produce soluble and insoluble factors critical for BA functions and homeostasis; conversely, BAs produce a range of growth factors and cytokines that collectively modulate vascular growth, survival, remodeling, regression, and blood perfusion [64,66,74]. In obese mice, the capillary density in BAT decreases significantly (i.e., ∼0.5 capillaries per BA) (**Figure 4**), resulting in hypoxia and BAT degeneration[65]. Our results showed that transplanted BA microtissues prevented the whitening of endogenous BAT (**Figure 4**), which may contribute to the improvement of glucose and insulin homeostasis. In addition, we showed transplanted BAs secreted human adipokines (**Figure 6A**) and altered the expression levels of many endogenous adipokines (**Fig 6B, 6C**). Thus, the transplanted microtissues function at least partially via the endocrine mechanism.

We found that the transplanted microtissues gradually became white adipocyte-like in morphology (**Figure 2B**), although UCP-1 proteins are still expressed at a high level (**Figure 2C**). A reason for this whitening is the insufficient vascularization in the transplant. This agrees well with literature findings that the transplants had fewer blood vessels than endogenous BAT [18,31]. Consequently, BAs gradually decreased UCP-1 expression and became WAT-like features (i.e., whitening). There are two potential ways to address this problem. First, as shown in a published study[18], supplementing vascular endothelial growth factor (VEGF) to the transplant can significantly improve its engraftment, angiogenesis, and function. Second, endothelial cells can be included in the BA microtissues. As stated in the above paragraph, endothelial cells or vasculature are indispensable for BAT in vivo.

We used immortalized BAPs to prepare mature BA microtissues. No tumor or abnormal tissue growth was observed (**Figure 2**), indicating fully differentiated BAs are safe in vivo. Thus, it is applicable to isolate and immortalize BAPs from a patient and expand them to prepare BA microtissues for personalized BA augmentation therapy. An alternative approach is to prepare induced pluripotent stem cells (hiPSCs) for a patient. hiPSCs can be generated by reprogramming somatic cells[75–77]. They have unlimited proliferation capability and can be differentiated into all types of somatic cells. Additionally, the transplant’s immune rejection can be avoided by preparing BAT from personalized hiPSCs[78,79]. Recent clinical studies showed autologous hiPSCs-derived dopaminergic neurons, and retinal cells were safe and effective in treating Parkinson’s[80] and macular degeneration[79], respectively, indicating the coming of the hiPSCs-based personalized medicine era. Alternatively, universal hiPSCs can be engineered, for instance, via inactivating major histocompatibility complex (MHC) class I and II genes and overexpressing CD47 or/and PD-L1[80–83]. The derivatives of universal hiPSCs are hypoimmunogenic and can be prepared at large scales as “off-the-shelf” allogeneic products. hiPSCs have been successfully differentiated into BAs that are metabolically active *in vitro* and in mice models[34,84–89]. Therefore, future studies can explore using hiPSCs to prepare personalized BA microtissues.

## Conclusion

In summary, our study showed that 3D BA microtissues could be fabricated at large scales, cryopreserved for the long term, and delivered via injection. BAs in the microtissues had higher purity, and higher UCP-1 protein expression than BAs prepared via 2D culture. In addition, 3D BA microtissues had good in vivo survival and tissue integration and had no uncontrolled tissue overgrowth. Furthermore, they showed good efficacy in preventing OB and T2DM with a very low dosage compared to literature studies. Thus, our results show engineered 3D BA microtissues are promising anti-OB/T2DM therapeutics. They have considerable advantages over dissociated BAs or BAPs for future clinical applications in terms of product scalability, storage, purity, quality, and in vivo safety, dosage, survival, integration, and efficacy.

## Abbreviations

OB: obesity
T2DM: type 2 diabetes mellitus
T1DM: type 1 diabetes mellitus
BAT: brown adipose tissue
BA: brown adipocyte
OCR: oxygen consumption rate
WAT: white adipose tissue
scWAT: subcutaneous white adipose tissue
STZ: streptozotocin
BAPs: brown adipocyte progenitors
UCP-1: uncoupling protein 1
ECM: extracellular matrix
WT: wide type mouse. Rag1^-/-^Rag 1 knock out mouse
HFD: high fat diet
NCD: normal chow diet
2D: two dimensional
3D: three dimensional
GTT: glucose tolerance test
ITT: Insulin tolerance test
endoBAT: endogenous BAT
MFI: mean fluorescent intensity

## Acknowledgments

This research was supported by the Nebraska DHHS Stem Cell Grant 2019 (YL), the Pennsylvania State University start-up (YL), and the University of Nebraska Collaboration Initiative Grant (YL, BD, SC).

## Competing financial interests

Authors have no competing financial interests.

**Supplementary information** accompanies this paper.

