## Supplemental figures for "Engineered human brown adipocyte microtissues improved glucose and insulin homeostasis in high fat diet-induced obese and diabetic mice"

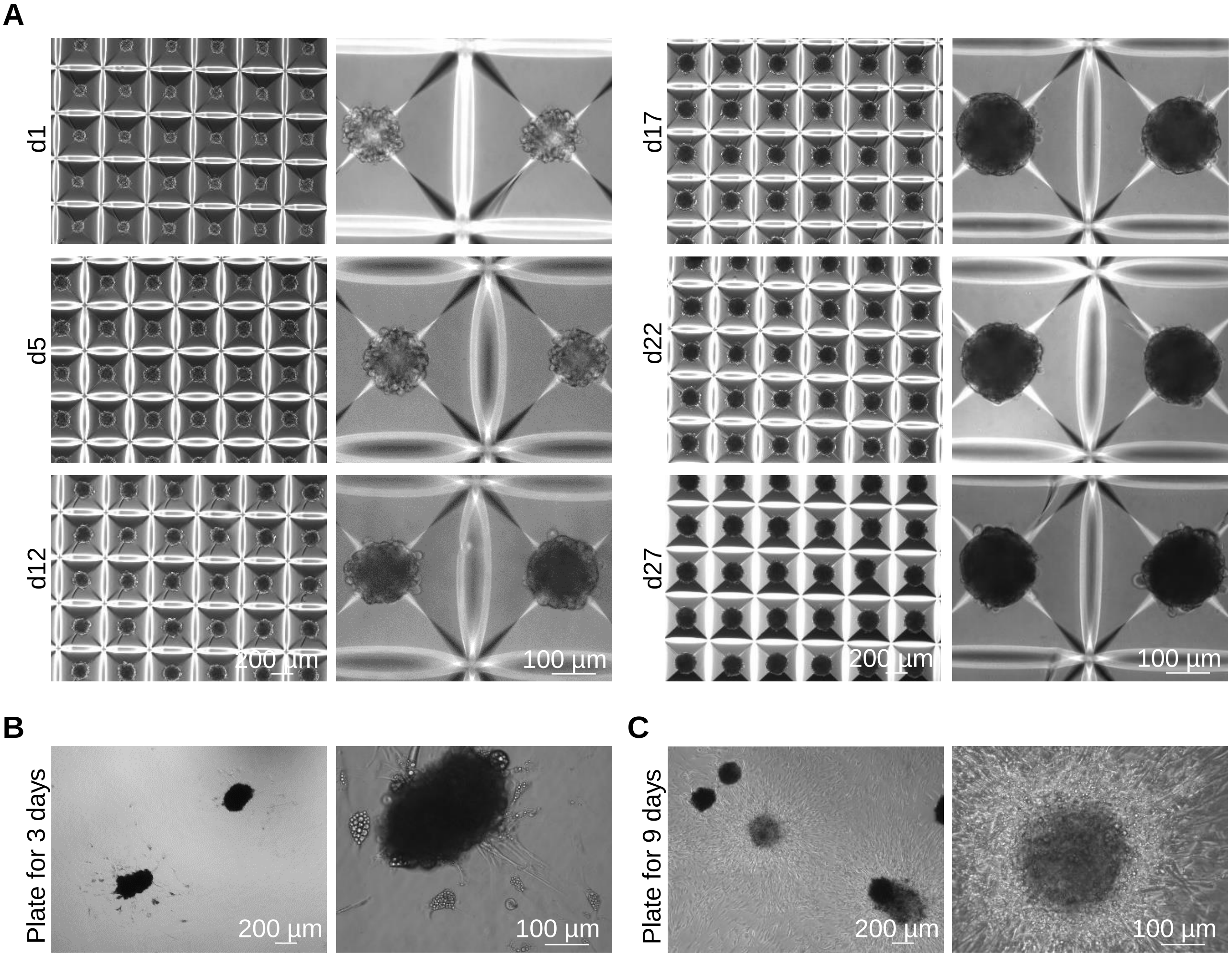


**Figure S1.** Fabricating BA microtissues. (**A**) Preparing BA microtissues in microwells. Phase images day-27 BA microtissues after plating on 2D surface for 3 days (**B**) and 9 days (**C**).

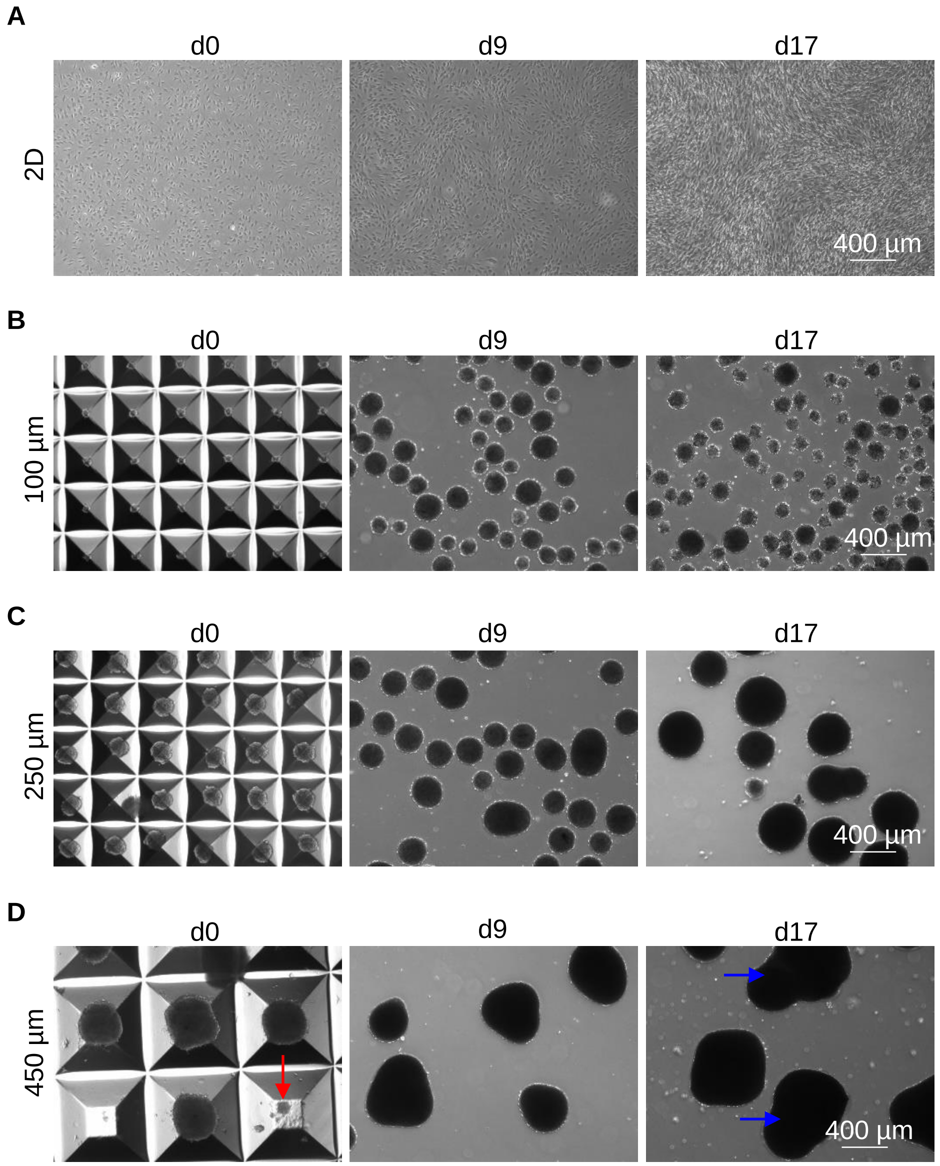


**Figure S2**. Preparing BAs in 2D culture (**A**) or microwells with varied aggregate sizes (**B, C, D**). Day 9 and Day 17 microtissues were released from microwells before imaging.

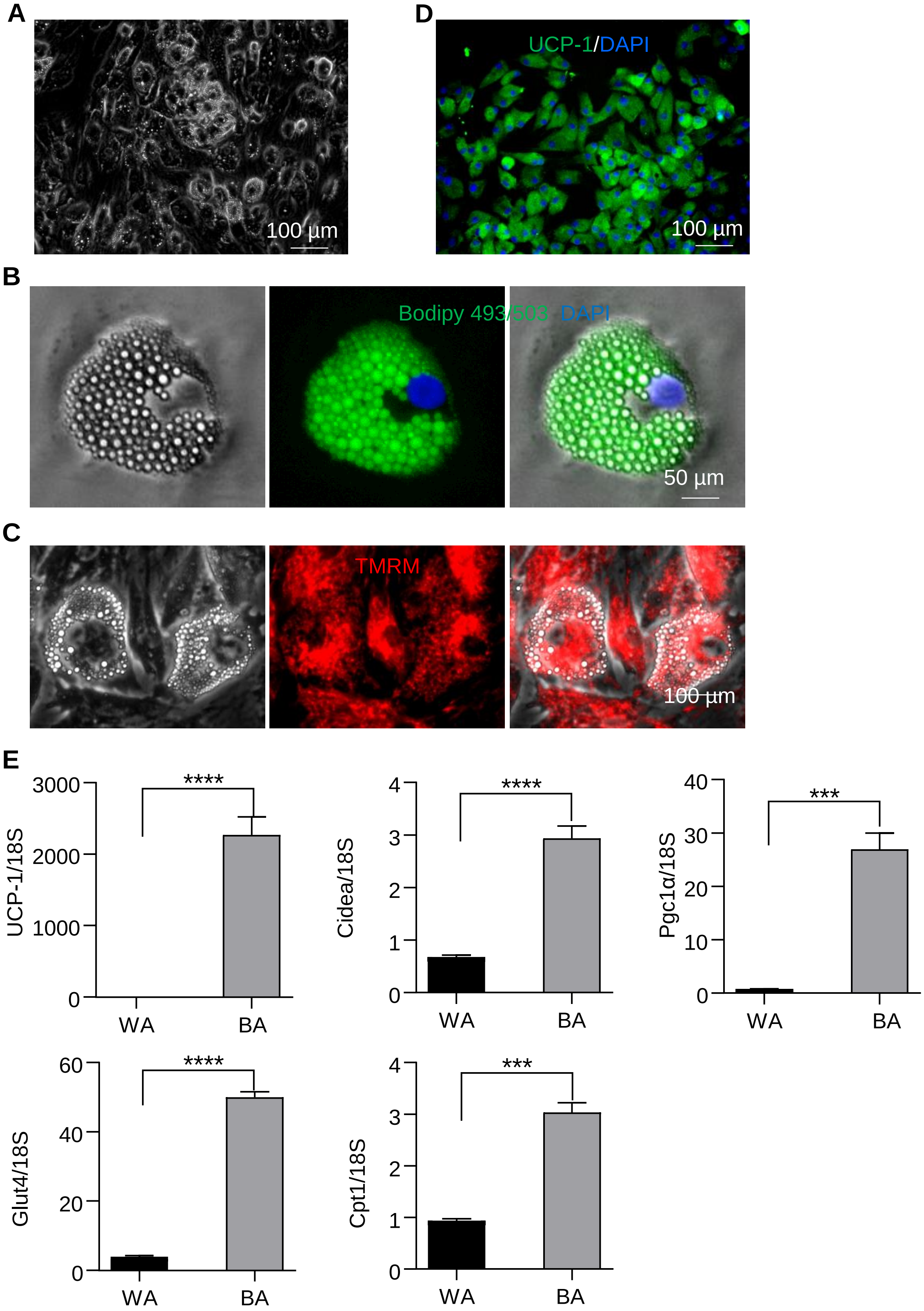


**Figure S3.** Characterization of BA microtissues. BAs prepared in 3D had typical BA phenotypes such as large numbers of small lipid droplets (**A, B**), abundant mitochondria (**C**) and UCP-1 proteins (**D**). They expressed BA-specific genes at high level (**E**). WA and BA: adipocytes differentiated from human WAPs and BAPs in microwells. TMRM: Tetramethylrhodamine, methyl ester (mitochondrial probe). Data are represented as mean ± SEM (n=3). ***p < 0.001, ****p < 0.0001 ***p < 0.001

**Table S1.** Antibodies used in this study.

| Antibody | Supplier | Catalog. No | Host species | Dilution |
|---|---|---|---|---|
| Anti-UCP1 | abcam | ab155117 | Rabbit | 1:50 |
| Tyrosine Hydroxylase (TH) | Fisher scientific | AB152MI | Rabbit | 1:50 |
| CD31 | abcam | ab24590 | Mouse | 1:250 |
| Anti-UCP1 | abcam | ab23841 | Rabbit | 1:250 |
| Anti-Human Nuclear Antigen antibody [235-1] | abcam | ab191181 | Mouse | 1:100 |
| Secondary Antibody | Thermo Fisher | A-21202 | Donkey | 1:500 |
| Secondary Antibody | Thermo Fisher | A-21207 | Donkey | 1:500 |
| Secondary Antibody | Jackson Immuno Research Labs | 715585150 | Donkey | 1:500 |
| Secondary Antibody | Jackson Immuno Research Labs | 711545152 | Donkey | 1:500 |

**Table S2.** One-way ANOVA multiple comparisons test results of mean UCP-1 intensities for day 17 BA in Figure 1D. *p < 0.05 , ****p < 0.0001.

| | Control | 2D BA | 100 µm | 250 µm | 450 µm |
|---|---|---|---|---|---|
| Control | | | | | |
| 2D BA | **** | | | | |
| 100 µm | **** | **** | | | |
| 250 µm | **** | **** | **** | | |
| 450 µm | **** | **** | **** | * | |

**Table S3.** Two-way ANOVA multiple comparisons test results of body weight gain (**A**), fat mass (**B**), lean mass (**C**), fasting glucose (**D**), GTT (**E**) and ITT (**F**) in Figure 3. *p < 0.05, **p < 0.01 , ***p < 0.001, ****p < 0.0001.

**A**

| Tukey's multiple comparisons test (weight) | WT NCD | WT HFD | $Rag1^{-/-}$NCD | $Rag1^{-/-}$HFD | $Rag1^{-/-}$ HFD+BAT |
|---|---|---|---|---|---|
| WT NCD | | | | | |
| WT HFD | **** | | | | |
| $Rag1^{-/-}$NCD | ns | **** | | | |
| $Rag1^{-/-}$HFD | **** | **** | **** | | |
| $Rag1^{-/-}$HFD+BAT | **** | ns | **** | **** | |

**B**

| Tukey's multiple comparisons test (fat mass) | WT NCD | WT HFD | $Rag1^{-/-}$NCD | $Rag1^{-/-}$HFD | $Rag1^{-/-}$ HFD+BAT |
|---|---|---|---|---|---|
| WT NCD | | | | | |
| WT HFD | **** | | | | |
| $Rag1^{-/-}$NCD | ns | **** | | | |
| $Rag1^{-/-}$HFD | **** | **** | **** | | |
| $Rag1^{-/-}$HFD+BAT | **** | ns | **** | **** | |

**C**

| Tukey's multiple comparisons test (lean mass) | WT NCD | WT HFD | $Rag1^{-/-}$NCD | $Rag1^{-/-}$HFD | $Rag1^{-/-}$ HFD+BAT |
|---|---|---|---|---|---|
| WT NCD | | | | | |
| WT HFD | **** | | | | |
| $Rag1^{-/-}$NCD | ns | **** | | | |
| $Rag1^{-/-}$HFD | **** | *** | **** | | |
| $Rag1^{-/-}$HFD+BAT | **** | ns | **** | *** | |

**D**

| Tukey's multiple comparisons test (fasting glucose) | WT NCD | WT HFD | $Rag1^{-/-}$NCD | $Rag1^{-/-}$HFD | $Rag1^{-/-}$ HFD+BAT |
|---|---|---|---|---|---|
| WT NCD | | | | | |
| WT HFD | ns | | | | |
| $Rag1^{-/-}$NCD | ns | ns | | | |
| $Rag1^{-/-}$HFD | **** | ** | **** | | |
| $Rag1^{-/-}$HFD+BAT | ** | ns | ns | * | |

**E**

| Tukey's multiple comparisons test (GTT) | WT NCD | WT HFD | $Rag1^{-/-}$NCD | $Rag1^{-/-}$HFD | $Rag1^{-/-}$ HFD+BAT |
|---|---|---|---|---|---|
| WT NCD | | | | | |
| WT HFD | ns | | | | |
| $Rag1^{-/-}$NCD | ns | ns | | | |
| $Rag1^{-/-}$HFD | **** | **** | **** | | |
| $Rag1^{-/-}$HFD+BAT | ** | ns | ns | **** | |

**F**

| Tukey's multiple comparisons test (ITT) | WT NCD | WT HFD | $Rag1^{-/-}$NCD | $Rag1^{-/-}$HFD | $Rag1^{-/-}$ HFD+BAT |
|---|---|---|---|---|---|
| WT NCD | | | | | |
| WT HFD | ns | | | | |
| $Rag1^{-/-}$NCD | ns | ns | | | |
| $Rag1^{-/-}$HFD | * | ns | *** | | |
| $Rag1^{-/-}$HFD+BAT | ns | ns | ns | * | |

**Table S4.** Two-way ANOVA multiple comparisons test results of mouse adipokine antibody array in Figure 6C. *p < 0.05, **p < 0.01, ***p < 0.001, ****p < 0.0001.

| | $Rag1^{-/-}$NCD vs. $Rag1^{-/-}$ HFD | $Rag1^{-/-}$NCD vs. $Rag1^{-/-}$ HFD+BA | $Rag1^{-/-}$HFD vs. $Rag1^{-/-}$ HFD+BAT |
|---|---|---|---|
| Adiponectin | **** | ns | ** |
| ANGPT-L3 | * | ns | * |
| C-Reactive Protein | * | ns | *** |
| ICAM-1 | * | ns | *** |
| IGF-I | *** | ns | *** |
| IGFBP-3 | **** | ns | **** |
| IGFBP-5 | ns | ns | * |
| IGFBP-6 | * | ns | * |
| Lipocalin-2 | ** | ns | ** |
| Pentraxin 2 | ns | ns | * |
